# Language and cognitive impairment associated to a novel p.Cys63Arg change in the MED13L gene

**DOI:** 10.1101/151688

**Authors:** M^a^ Salud Jiménez-Romero, Pilar Carrasco-Salas, Antonio Benítez-Burraco

## Abstract

Mutations of the *MED13L* gene, which encodes a subunit of a transcriptional regulatory complex, result in a complex phenotype entailing physical and cognitive anomalies. Deep language impairment has been reported, mostly in patients with CNV. *Case presentation.* We report on a child who presents with a non-synonymous change p.Cys63Arg in *MED13L* (Chr12:116675396A>G, GRCh37) and who exhibits profound language impairment in the expressive domain, cognitive delay, behavioral disturbances, and some autistic features. *Conclusions.* Because of the brain areas in which *MED13L* is expressed and because of the functional links between MED13L and the products of some candidate genes for language disorders, the proband’s linguistic phenotype may result from changes in a functional network important for language development.

## INTRODUCTION

Rare conditions involving language deficits and resulting from point mutations in the coding region of genes provide crucial evidence of how the linguistic brain is wired in response to genetic guidance. The *MED13L* gene encodes a subunit of the mediator complex (a scaffolding protein device that promotes the assembly of functional preinitiation complexes with RNA polymerase II and the general transcription factors) that seemingly helps regulate the transcription of targets of the Wnt, Notch, and Shh signalling pathways (Asadollahi et al. 2013, Davis et al. 2013, Utami et al. 2014). Among other tissues, *MED13L* gene is highly expressed in the fetal and adult brain, particularly in the cerebellum after birth (Muncke et al. 2003, Musante et al. 2004; Human Brain Transcriptome: http://hbatlas.org/hbtd/images/wholeBrain/MED13L.pdf). *MED13L* alteration was initially reported to cause complex heart defects, but it is now clear that mutations of *MED13L* appear to underlie a gradient, also giving rise to facial dysmorphisms, intellectual disability, delayed speech and language, hypotonia, and/or behavioral difficulties (Adegbola et al. 2015, Cafiero et al. 2015). It has been hypothesized that these diverse effects may result from the involvement of the gene in neural crest induction (see Asadollahi et al. 2013 for discussion). As noted by Cafiero et al (2015), *MED13L* haploinsufficiency, resulting from CNVs or from loss-of-function intragenic variants, causes a recognizable intellectual disability (ID) syndrome with distinctive facial features and with or without heart defects. Nevertheless, the clinical relevance of missense mutations in the *MED13L* gene is poorly understood, and only some cases have been described to present. This is particularly true regarding their effect on language and cognition.

In this paper, we report on a boy with a missense mutation affecting the N-terminal region of the MED13L protein, and provide a detailed characterization of his clinical features, with a focus on the linguistic and cognitive phenotype. We discuss our findings in the context of current studies of the genetic basis of language.

## METHODS

### Linguistic, cognitive, and behavioral assessment

Global development of the boy was assessed with the Spanish version of the Battelle Developmental Inventories (De la Cruz and González, 2011). This test comprises 341 items and evaluates gross and fine motor skills, adaptive abilities, personal/social development, cognitive development, and receptive and expressive communication skills.

The expressive and communicative abilities of the child were assessed in more detail with the Spanish version of the MacArthur-Bates Communicative Development Inventories (López-Ornat, 2005). This is a parent report instrument, which aims to characterize early language development. The test evaluates several areas, including vocalizations, first words, gestures, vocabulary comprehension, and grammar.

The perceptual and understanding abilities of the boy were also assessed with the Spanish version of the ComFor Test (Verpoorten et al. 2014). This test measures perception and sense-making at the levels of presentation and representation, with the aim of designing accurate augmentative communication interventions for children with profound language impairment. It encompasses 36 items, arranged in five series and two levels. Level I comprises series 1 to 3 and evaluates perception: objects or pictures have to be sorted according to perceptible features (shape, colour, matter and size). Level II comprises series 4 and 5 and evaluates representation: objects or pictures have to be sorted on the basis of non-perceptible features.

Finally, the Spanish version of the Modified Checklist for Autism in Toddlers (M-CHAT) (Robins et al. 2001) was used for screening autistic features in the child. This test is designed for toddlers between 18 and 60 months and comprises 23 questions. A positive score is indicative of the possibility of suffering from the disease.

### Molecular and cytogenetic analysis

DNA from the patient and his parents was extracted from 100 μl of EDTA-anticoagulated whole blood using MagNA Pure (Roche Diagnostics, West Sussex, UK) and used for subsequent analyses.

#### Fragile X syndrome determinants

CGG expansions affecting the gene *FMR1* (the main determinant for X-fragile syndrome) were analyzed in the patient according to standard protocols. Polymerase chain reaction (PCR) of the fragile site was performed with specific primers and trinucleotide repeat size of the resulting fragments were evaluated.

#### Microrrays for CNVs search and chromosome aberrations analysis

Patient DNA was hybridized on a CGH platform (Cytochip Oligo ISCA 60K). The DLRS value was > 0.10. The platform included 60.000 probes. Data were analyzed with Agilent Genomic Workbench 7.0 and the ADM-2 algorithm (threshold = 6.0; aberrant regions had more than 5 consecutive probes).

#### Whole-exome sequencing (WES)

The panel used for the preparation of the library was designed by SureSelectXT Human All Exon V5 (Agilent Technologies). It captures > 35000 exons from > 19000 genes and the splicing flanking (5 bp) regions. The size is ~50Mb. Sequencing was performed with the HiSeq 2500 SystemTM (Illumina) sequencer. The reads obtained were filtered, based on quality parameters, and aligned to the reference genome (build 37 of genome Hg19), using the BWA alignment program (version 0.7.12). Sequence variations with respect to reference genome were detected using the GATK algorithm (version 2.3-9). Variants were annotated with the latest available version of ION Reporter (Life Technologies) and an in-house developed pipeline, aimed to register the patient’s variants according to a genetic type classification. Our analysis focused on variants found in coding or splicing regions. Changes included missense, nonsense and stoploss mutations, as well as nucleotide insertions or deletions, found in > 40% of the sequences obtained.

#### Sanger sequencing

For analysing in the patient’s parents the variants of unknown significance identified by the WES study of the proband, a PCR was performed using specific primers for exons 2 and 37 of *MED13L* and *ANK3* genes, respectively, according to the reference sequences NM___15335 and NM020987. Double-sense sequencing of PCR products was done in an automatic sequencer ABI 3130 DNA Analyzer (Applied Biosystems, Foster City, California, USA), with further study of sequence variants using software seqScape v2.5 (Life Technologies, CA).

## RESULTS

### Clinical History

The proband is a boy born after 38 weeks of normal gestation by eutocic deliver to a healthy 37 years old female. The most relevant clinical data at birth are summarized in table 1.

**Table 1.**
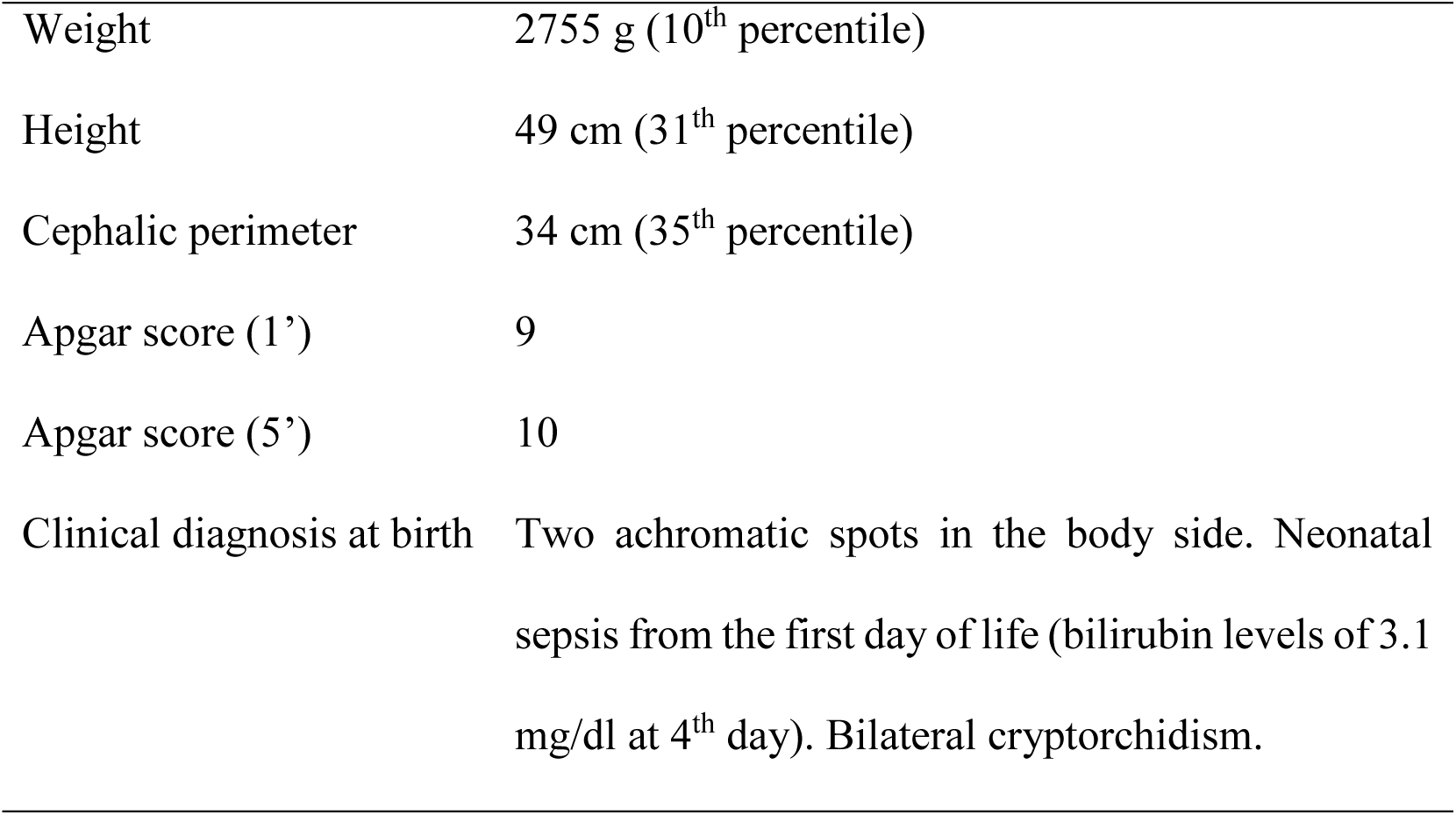
Summary table with the most relevant clinical findings at birth.

|  |  |
| --- | --- |
| Weight | 2755 g (10<sup>th</sup> percentile) |
| Height | 49 cm (31<sup>th</sup> percentile) |
| Cephalic perimeter | 34 cm (35<sup>th</sup> percentile) |
| Apgar score (1') | 9 |
| Apgar score (5') | 10 |
| Clinical diagnosis at birth | Two achromatic spots in the body side. Neonatal sepsis from the first day of life (bilirubin levels of 3.1 mg/dl at 4<sup>th</sup> day). Bilateral cryptorchidism. |

No signs of cardiac disease were observed. Auricular and ventricular volumes were concordant and no abnormal signs of dilatation were found. Auriculoventricular and sigmoid valves showed normal morphology and kinetics, and no cardiac insufficiency was detected. Global and segmental ventricular contraction was normal. The patient exhibited a type I septum IV and a mild, non-pathological hypertrabeculation in the apical zone of the left ventricle. M-Mode measurements were as follows: Ejection Fraction: 61%; interventricular septum end-diastolic thickness: 6mm; left ventricular end-diastolic diameter: 33mm; left ventricular end-systolic diameter: 22.5mm; left-ventricular posterior wall thickness: 5.6mm.

Cranial magnetic resonance imaging (MRI), performed at age 3;5 (years;months), yielded normal results. An electroencephalography (EEG) study with sleep deprivation was performed at age 4;9. and revealed frequent epileptiform discharges during sleep in the left parietotemporal region and right centrotemporal region in absence of continuous spikes and waves during slow wave sleep.

Developmental anomalies were soon reported by the parents, including lack of crying, absence of feeding demands, sucking weakness, reduced eye contact, hypotonia, and neck rigidity. Still, the baby seemed to enjoy physical contact and smiled regularly. Motor development was also delayed. Generalized hypotonia was confirmed at 6 months. At age 2;0 mild axial hypotonia was still observed, whereas at age 4;10 some hypertonic symptoms were found by the neuropediatrician. Self-hurting behavior was reported at age 4;10. The boy only started to walk without aid at age 3;0. Stereotypic movements were reported from age 3;0. Dysmorphic features included reduced palpebral fissures, nasal base enlargement, enlarged plane philtrum, a thinner upper lip, low-set ears, and a prominent columella. Hypermetropia was diagnosed at age 4;10.

The boy started attending a normal nursery school at age 1;4, but a month later was transferred to an early care center for disabled children, where he was reported to interact with his peers normally. At present (age 5;4) he is attending a normal school. Self-hurting behavior is still present. Sphincter control has not been yet achieved. Feeding problems are still observed, to the extent that he is only able to eat grinded food.

### Language and cognitive development

Early language milestones were achieved late by the child. Cooing was first observed at six months, whereas reduplicated babbling was observed at age 1;6 only. Learning of a first word (*agua* ‘water’) was reported at age 1;0, but it was lost soon and no further progress was observed in his expressive abilities. At age 2;0 the boy didn’t attend others upon request and was unable to understand simple commands.

From age 1;5 onwards the child started to received speech therapy 4 times a week. Comprehension improved and at present, he is able to understand forbidding utterances (‘Don’t do this’, ‘Don’t touch that’) and usually reacts to simple direct commands (‘Come here!’, ‘Give me that’). Nonetheless, his expressive abilities remain at a prelinguistic stage. He still babbles, particularly when he is playing alone or before going to sleep. These nonsense disyllabic sequences have no real communicative function and resemble those generated during the canonical babbling stage. Still, some communicative intentionality is evident in his behavior. For instance, he uses to hold the other’s hand to lead him/her to some object of interest. Nonetheless, deictical pointing has not been reported. Also, no signs of symbolic behavior have been observed, including differed imitation, drawing, or playing with objects simulating animals or persons. Echolalias are absent, but hypersensibility to loud, unfamiliar sounds has been reported by the parents. At age 4;10 the neuropediatrician diagnosed the child as suffering from syndromic autism spectrum disorder (ASD) with no language and with mental retardation. However, the boy scored 2 points in the M-Chat test, which is indicative of a low risk for the disease. Still, some autistic behaviors have been reported by his parents. Specifically, the boy does not use his index finger to point or follow other’s pointing, he makes unusual finger movements near his eyes, he does not play pretend or make-believe, and he exhibits restricted interests and stereotyped behaviors.

We have assessed in depth his global development at age 5;4 with the Spanish version of the Battelle Developmental Inventories. The resulting scores are suggestive of a broad developmental delay, mostly impacting on his language abilities. Accordingly, communication skills are severely impaired; adaptive and personal-social abilities are impaired; and motor and cognitive skills are the less affected areas (figure 1).

**Figure 1.**
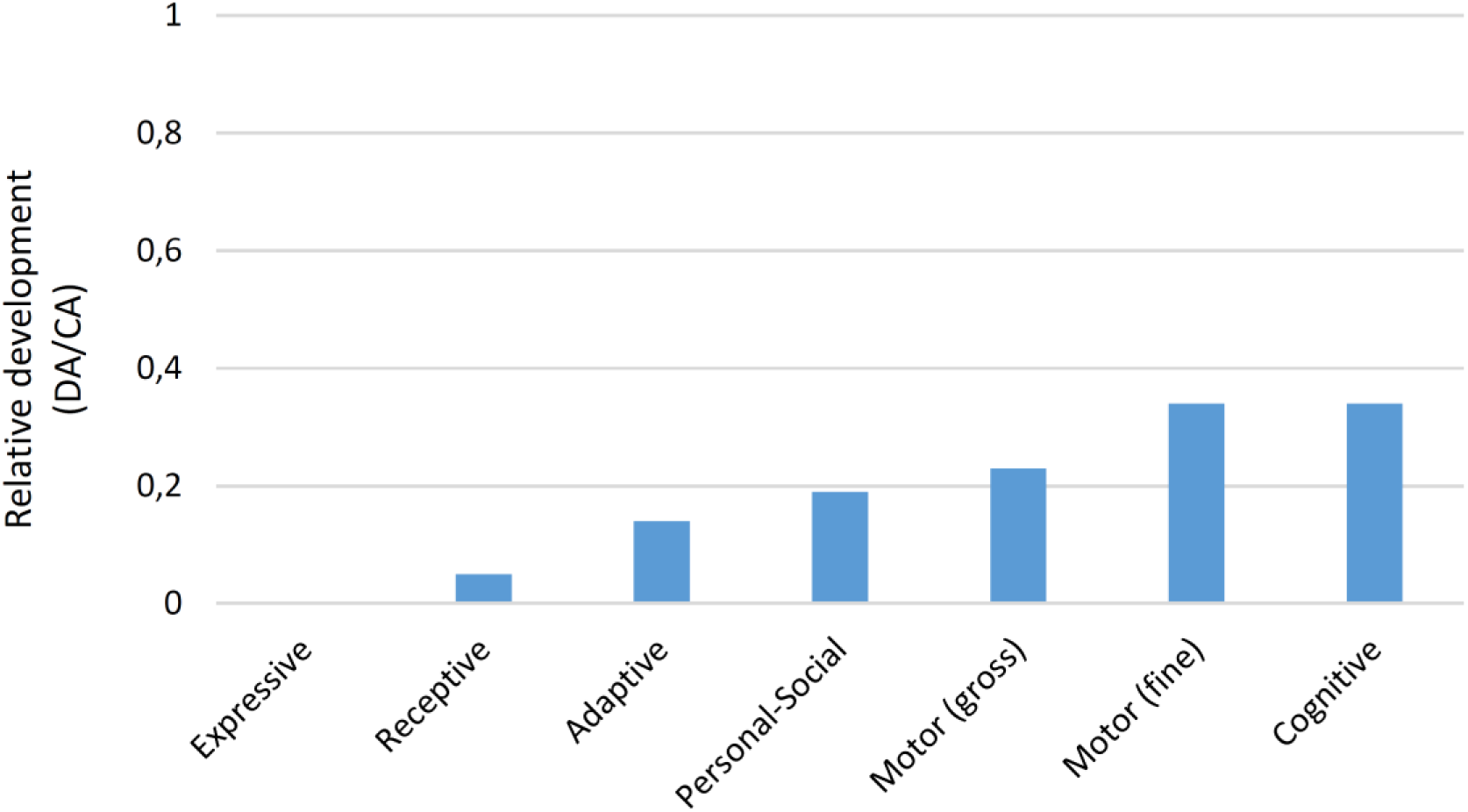
The proband’s developmental profile at age 5;4 according to the Battelle Developmental Inventories. In order to make reliable comparisons, the resulting scores are shown as relative values referred to the expected scores according to the chronological age of the child. Abbreviations: DA, developmental age; CA, chronological age.

Additionally, we evaluated his communicative abilities with the Spanish version of the MacArthur-Bates Communicative Development Inventories. The obtained scores suggest that the boy is still at a prelinguistic stage. Regarding his vocal behaviour, some milestones have been already achieved (vocal playing is frequently elicited by music listening, babbling is used for requesting, and mimicry of the prosodic envelope of requests or questions have been reported).

In the ComFor test, which evaluates precursors of communication ability, he performed correctly series 1 to 3 from Level I, but failed with Level II tasks., suggesting that his representation abilities are severely affected.

### Molecular and cytogenetic analysis

Routine molecular and cytogenetic analyses of the boy were performed at ages 2;7-2;8. Polymerase chain reaction (PCR) analysis of the FMR1 fragile site was normal. Patient has a normal allele of 30 CGG repeats. No chromosomal rearrangements nor CNVs were observed in the comparative genomic hybridization array (array-CGH). The whole-exome sequencing (WES) analysis revealed several variants in genes that are associated with conditions that share some or most of the symptoms and features exhibited by our proband (table 2). Considering the clinical profile of the subject, the most promising variant is the c.187T>C transition (NM_015335.4) in heterozygous state in exon 2 of the *MED13L* gene (Figure 2A), resulting in the change p.Cys63Arg in the Med13_N terminal domain of the protein (http://pfam.xfam.org/protein/Q71F56) (figure 2B). This cysteine residue is conserved across all primates, excepting bushbabies and tarsiers, and is found in >70% of bony vertebrates (figure 2C). This is a novel variant not previously reported in the public databases (dbSNP, ExAC, HGMD, LOVD and NHLBI ESP). The segregation study performed in the patient’s parents revealed that this is a de novo substitution. In addition to this probably pathogenic mutation, two heterozygous missense changes in exon 37 of the *ANK3* gene, reported with low frequency in ExAC database, were identified: c.9997A>T (p.Thr3333Ser) and c.10688A>G (p.Glu3563Gly) (NM_020987.3). These variants in the *ANK3* gene have been inherited from his asymptomatic mother and therefore, were reclassified as probably benign variants. Finally, the WES analysis revealed five other low frequency variants (< 1% in the general population) of potential interest: c.4282T>C (p.Ser1428Pro) (NM_015338.5) in *ASXL1*, c.1733C>T (p.Ser578Phe) (NM_030632.1) in *ASXL3*, c.4855A>C (p.Lys1619Gln) (NM_033427.2) in *CTTNBP2*, c.11380C>T (p.Pro3794Ser) (NM_033427.2) in *KMT2D*, and c.529G>A (p.Ala177Thr) (NM_024570.3) in *RNASEH2B*. These genes are involved in brain development and function and their mutation give rise to some of the symptoms or features observed in our proband. For example, both *ASXL1* and *ASXL3* are candidates for Bohring-Opitz syndrome (#605039) and Bainbridge-Ropers syndrome (#615485), both with autosomal dominant inheritance and both entailing intellectual disability and dysmorphic features (Shashi et al. 2016). However, the clinical profile of the patient and the nature of these variants leaded us to reclassify them as not relevant for genotype-to-phenotype correlations.

**Table 2.**
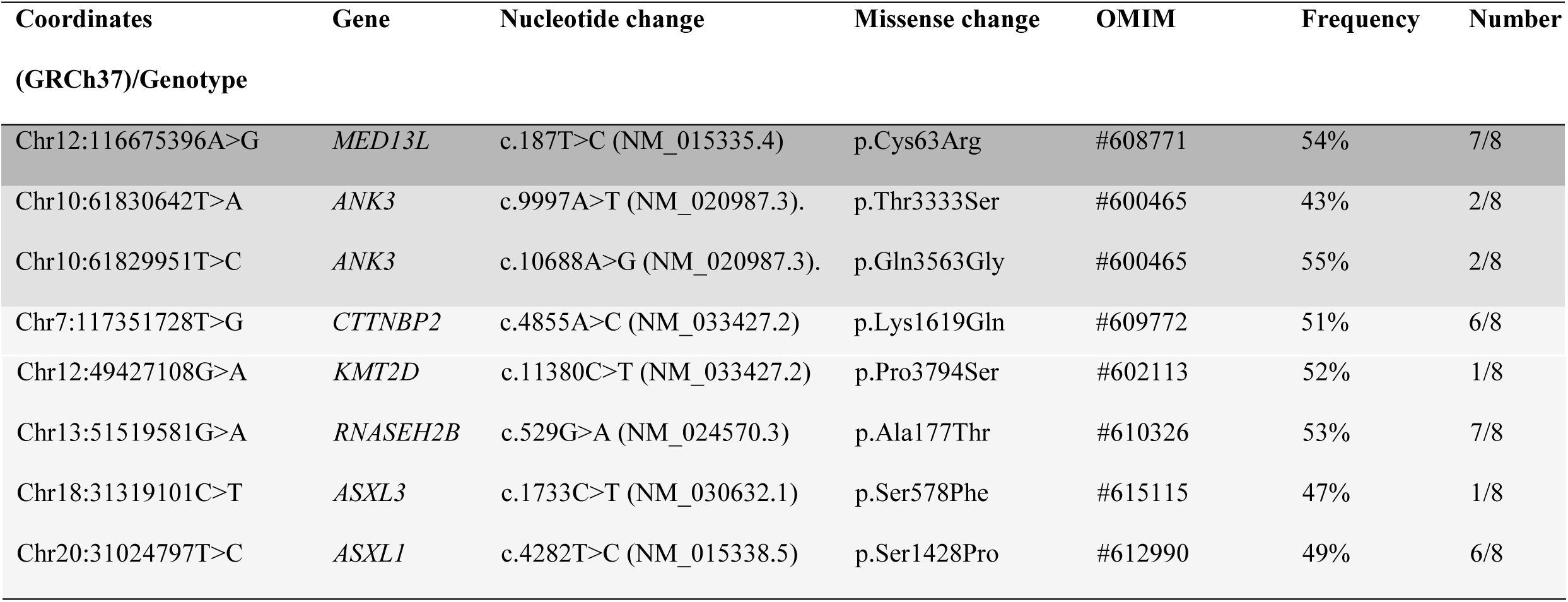
Missense mutations as revealed by the WES analysis of the proband. Mutations of clinical interest are shadowed in medium and dark grey.

| Coordinates<br>(GRCh37)/Genotype | Gene | Nucleotide change | Missense change | OMIM | Frequency | Number |
| --- | --- | --- | --- | --- | --- | --- |
| Chr12:116675396A>G | *MED13L* | c.187T>C (NM_015335.4) | p.Cys63Arg | #608771 | 54% | 7/8 |
| Chr10:61830642T>A | *ANK3* | c.9997A>T (NM_020987.3). | p.Thr3333Ser | #600465 | 43% | 2/8 |
| Chr10:61829951T>C | *ANK3* | c.10688A>G (NM_020987.3). | p.Gln3563Gly | #600465 | 55% | 2/8 |
| Chr7:117351728T>G | *CTTNBP2* | c.4855A>C (NM_033427.2) | p.Lys1619Gln | #609772 | 51% | 6/8 |
| Chr12:49427108G>A | *KMT2D* | c.11380C>T (NM_033427.2) | p.Pro3794Ser | #602113 | 52% | 1/8 |
| Chr13:51519581G>A | *RNASEH2B* | c.529G>A (NM_024570.3) | p.Ala177Thr | #610326 | 53% | 7/8 |
| Chr18:31319101C>T | *ASXL3* | c.1733C>T (NM_030632.1) | p.Ser578Phe | #615115 | 47% | 1/8 |
| Chr20:31024797T>C | *ASXL1* | c.4282T>C (NM_015338.5) | p.Ser1428Pro | #612990 | 49% | 6/8 |

**Figure 2.**
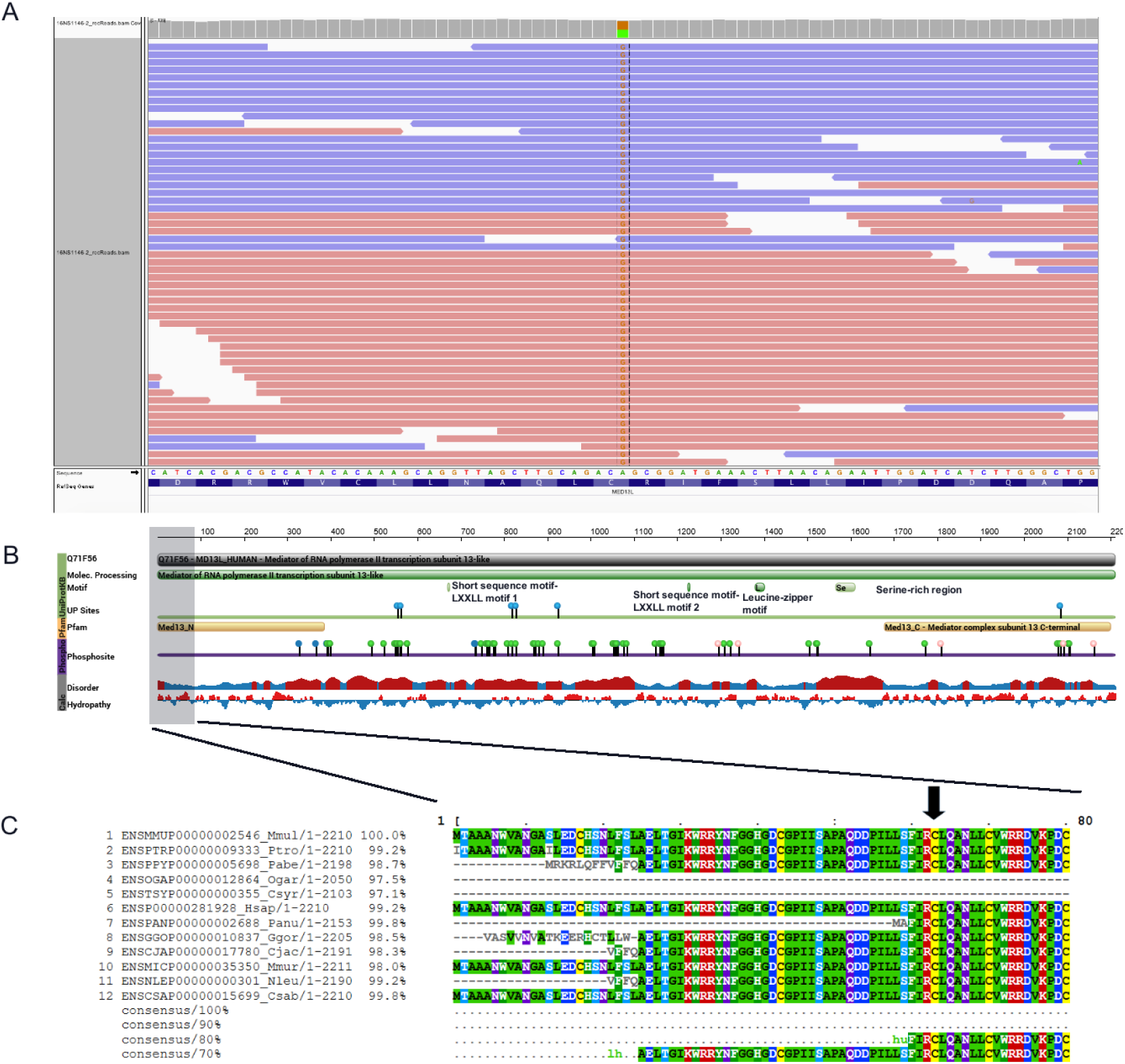
Molecular aspects of the MED13L alteration in our proband. A. Outcome of the WES analysis of the patient showing the missense change in position Chr12:116237591. B. Main structural features of the MED13L protein. The scheme was obtained from the Protein Feature View tool of the Protein Data Bank (PDB) (http://www.rcsb.org/pdb/protein/Q71F56?evtc=Suggest&evta=ProteinFeatureView&evtl=autosearch_SearchBar_querySuggest). The vertical color bar on the left side indicates data provenance. Data in green originates from UniProtKB. Data in yellow originates from Pfam, by interacting with the HMMER3 web site. Data in purple originates from Phosphosite. Data in grey has been calculated using BioJava. Protein disorder predictions are based on JRONN (Troshin, P. and Barton, G. J. unpublished), a Java implementation of RONN (Red: potentially disordered region; Blue: probably ordered region). Hydropathy has been calculated using a sliding window of 15 residues and summing up scores from standard hydrophobicity tables (Red: hydrophobic; Blue: hydrophilic). C. Alignment of primate MED13L proteins using the MView tool of EMBL-EBI (http://www.ebi.ac.uk/Tools/msa/mview/). Conserved aminoacids are displayed at the bottom. The black arrow at the top points to the Cys changed into Arg in our proband.

### Discussion

The development of next generation sequencing facilities has allowed identify many new genes and many new variants of known genes related to clinical conditions entailing problems with language. In this paper, we have characterized the linguistic and cognitive profile of a boy with a novel heterozygous missense mutation in the *MED13L* gene, which is predicted to affect to the N-terminal domain of the protein. Our proband exhibits most of the cognitive and behavioral features of the *MED13L* haploinsufficiency syndrome, without congenital heart defect, and particularly, all the abnormal features linked to missense mutations in the gene (table 3), being language the most impaired domain. Accordingly, speech is severely affected and comprehension is limited to simple, direct commands. No alternative communication system is still available to the child and in fact, the results achieved in the ComFor test predict problems for acquiring an Augmentative and Alternative Communication (ACC). At present, language impairment related to MED13L dysfunction has been mostly reported in patients bearing point nonsense mutations of the gene (DECIPHER patients 258675 and 261065), frameshift mutations (DECIPHER patients 260083, 273003, and 323645), or partial deletions of *MED13L* (DECIPHER patients 283932 and 322949). It is thus of interest to know more about the effect on the language abilities of the affected children of point changes in the functional domains of MED13L.

**Table 3.** Summary table with the most relevant clinical features of our proband compared to other cases involving missense mutations of MED13L and to the common presentation of the MED13L haploinsufficiency syndrome.

|  | Cys63Arg<sup>1</sup> | Asp860Gly<sup>2</sup> | Pro866Leu<sup>3</sup> | Pro866Ala<sup>4</sup> | Cys1160Arg<sup>5</sup> | Arg1416His<sup>6</sup> | Gly1899Arg<sup>7</sup> | Gly1899Arg<sup>8</sup> | Ser2002Leu<sup>9</sup> | Thr2162Met<sup>10</sup> | Key phenotype features of MED13L-associated disease<sup>11</sup> |
| --- | --- | --- | --- | --- | --- | --- | --- | --- | --- | --- | --- |
| Cardiac anomalies | - | ? | ? | + | + | ? | ? | ? | ? | ? | + |
| Visual problems | + | ? | ? | + | ? | ? | ? | ? | + | ? |  |
| Facial dysmorphisms | + | + | + | + | ? | ? | + | + | + | + | + |
| Motor problems |  |  |  |  |  |  |  |  |  |  |  |
| Motor delay | + | ? | ? | ? |  | ? | ? | ? | + | ? |  |
| Hypotonia | + | ? | ? | ? | + | ? | ? | + | + | ? | + |
| Skull anomalies | - | ? | + | + | ? | ? | + | ? | ? | ? | + |
| Brain structural anomalies | - | ? | ? | + | ? | ? | ? | ? | + | ? |  |
| EEG anomalies | + | + | ? | + | + | ? | ? | ? | ? | ? |  |
| Cognitive impairment |  |  |  |  |  |  |  |  |  |  |  |
| Behavioural problems | + | ? | ? | ? | ? | ? | ? | + | ? | + | + |
| Developmental delay | + | ? | ? | + | + | ? | ? | ? | ? | ? |  |
| Intellectual disability | + | + | + | ? | ? | + | ? | + | + | + | + |
| Speech/language impairment | + | ? | ? | ? | ? | ? | ? | ? | ? | + |  |

1. Our proband
2. Trio 22 from Gilissen et al. 2014
3. DECIPHER patient 258131
4. DECIPHER patient 268019
5. DECIPHER patient 265953
6. Family M142 from Najmabadi et al. 2011
7. DECIPHER patient 262545
8. DECIPHER patient 323183
9. DECIPHER patient 260542
10. DECIPHER patient 272205
11. Summarized from Adegbola et al. 2015 (Table 3)
NB. The four missense mutations found by Muncke et al. (2003) (K686Q, E251G, R1872H, D2023G) are reported to entail cardiac problems only: no behavioural or cognitive deficits are mentioned in their paper)

*MED13L* is a candidate for ASD (Iossifov et al. 2012) and this could explain the ASD features of the proband, which includes language delay, no deictical pointing, absence of symbolic behavior, and stereotyped behavior. Additionally, MED13L is functionally related to several candidates for language and cognitive disorders, specifically with *EP300* and *CREBBP* (Krebs et al. 2010, Galbraith et al. 2013). Interestingly, the proteins encoded by these two genes serve as link between the FOXP2 and ROBO1 networks, which play a central role in the regulation of brain areas involved in speech and language processing (see Boeckx and Benítez-Burraco 2014 for details). Also, mutations in both EP300 and CREBBP are candidates for different subtypes of Rubinstein-Taybi syndrome (OMIM#613684 and OMIM#180849, respectively), a clinical condition entailing mental and growth retardation, skeletal abnormalities, and brain abnormalities mostly impacting in the rolandic area, which is crucial for language processing (Sener, 1995, Viosca et al. 2010). Interestingly too, Crebbp(+/-) mice exhibit altered vocal behavior (Wang et al. 2010) and the knock-out of the gene in postmitotic neurons of the forebrain results in problems for object recognition (Valor et al. 2011). Regarding to the *EP300* gene, mutations in the gene usually give rise to less severe mental impairment, more severe microcephaly, and more changes in facial bone structure (Hennekam, 2006). Finally, we wish highlight the role played by *MED13L* in neural crest induction. Interestingly, neurocristopathies (that is, conditions resulting from neural crest defects) usually entail skull, cognitive, and language features (see Benítez-Burraco et al. 2016 for discussion), but may also involve autistic features too, as observed in CHARGE syndrome (OMIM#214800) (Fernell et al. 1999). It is also worth highlighting the involvement of MED13L in the Shh pathway, which plays a key role in neural crest cell proliferation and fate (Wada et al. 2005, Calloni et al. 2007, Delloye-Bourgeois et al. 2014). The Shh pathway also regulates axonal growth and synaptogenesis in different parts of the brain (Bourikas et al. 2005; Salie et al. 2005). An excess of SHH activity results in facial dysmorphisms, mental retardation, and lack of language acquisition (Guion-Almeida and Richieri-Costa, 2009). Lastly, Shh also regulates the topographical sorting of thalamocortical axons and motor neurons important for speech (see Boeckx and Benítez-Burraco, 2014 for details).

## CONCLUSIONS

Although the exact molecular causes of the language and cognitive impairment exhibited by our proband remain to be elucidated, we expect that they result from the change we have uncovered in a conserved residue of the N-terminal domain of the MED13L protein. At the best of our knowledge, the missense mutation found in our proband is the second missense mutation in *MED13L* explicitly linked to language problems. In our case, we have provided with an exhaustive characterization of the language impairment observed in the child bearing this novel mutation. We have hypothesized that this mutation might have altered the functionality of the protein and/or its translocation to the nucleus, thus reducing the available amount of functional MED13L and impacting on the functions performed by the protein during embryonic development and after birth as part of the mediator complex. We expect that this clinical case helps improve our knowledge of the clinical presentation of missense variants within this important gene and ultimately, for our understanding of the genetic foundations of the human faculty for language.

## ETHICS, CONSENT, AND PERMISSIONS

Ethics approval for this research was granted by the Comité Ético del Hospital “Juan Ramón Jiménez”. Written informed consent was obtained from the probands’ parents for publication of this case report and of any accompanying tables and images. A copy of the written consent is available for review by the Editor-in-Chief of this journal.

## LIST OF ABBREVIATIONS

ACC: Augmentative and Alternative Communication
ASD: Autism Spectrum Disorder
CA: chronological age.
CGH: comparative genomic hybridization
CNVs: copy number variants
DA: developmental age
dbSNP: The Short Genetic Variations database (https://www.ncbi.nlm.n)
EEG: electroencephalography
ExAC: The Exome Aggregation Consortium (http://exac.broadinstitute.org/)
FISH: fluorescence in situ hybridization
HGMD: The Human Gene Mutation Database (http://www.hgmd.cf.ac.uk/ac/index.php)
LOVD: Leiden Open Variation Database (http://www.lovd.nl/3.0/home)
M-CHAT: Modified Checklist for Autism in Toddlers
MRI: magnetic resonance imaging
NHLBI ESP: the National Heart, Lung, and Blood Institute (NHLBI) GO Exome Sequencing Project (ESP) (http://evs.gs.washington.edu/EVS/)
PCR: polymerase chain reaction
WES: Whole-Exome Sequencing

## AUTHORS’ CONTRIBUTIONS

MJR assessed the cognitive and linguistic problems of the child, and made substantial contributions to the analysis of the data. MCS performed the molecular cytogenetic analyses. ABB interpreted the data and wrote the paper. All authors revised the draft of the paper, and read and approved the final manuscript.

## AUTHORS’ INFORMATION

M^a^ Salud Jiménez Romero is an Assistant Professor at the Department of Psychology of the University of Córdoba (Spain). M^a^ Pilar Carrasco Salas is a staff physician at the Human Genetics Unit of the Hospital “Juan Ramón Jiménez” in Huelva (Spain). Antonio Benítez-Burraco is Associate Professor at the Department of Philology of the University of Huelva (Spain).

## ACKNOWLEDGMENTS

We would like to thank the proband and his family for their participation in this research. Preparation of this work was supported by funds from the Spanish Ministry of Economy and Competitiveness (grant FFI2016-78034-C2-2-P to Antonio Benítez-Burraco).

## REFERENCES

Adegbola A, Musante L, Callewaert B, Maciel P, Hu H, Isidor B, Picker-Minh S, Le Caignec C, Delle Chiaie B, Vanakker O, Menten B, Dheedene A, Bockaert N, Roelens F, Decaestecker K, Silva J, Soares G, Lopes F, Najmabadi H, Kahrizi K, Cox GF, Angus SP, Staropoli JF, Fischer U, Suckow V, Bartsch O, Chess A, Ropers HH, Wienker TF, Hübner C, Kaindl AM, and Kalscheuer VM (2015). Redefining the MED13L syndrome. Eur J Hum Genet. 23:1308-17. doi: 10.1038/ejhg.2015.26.

Asadollahi R, Oneda B, Sheth F, Azzarello-Burri S, Baldinger R, Joset P, Latal B, Knirsch W, Desai S, Baumer A, Houge G, Andrieux J, and Rauch A. (2013) Dosage changes of MED13L further delineate its role in congenital heart defects and intellectual disability. Eur J Hum Genet. 21:1100-4. doi: 10.1038/ejhg.2013.17.

Benítez-Burraco A, Theofanopoulou C and Boeckx C (2016) Globularization and domestication. Topoi doi: 10.1007/s11245-016-9399-7

Boeckx C and Benítez-Burraco A (2014) Globularity and language-readiness: Generating new predictions by expanding the set of genes of interest. Front Psychol 5:1324. doi: 10.3389/fpsyg.2014.01324

Bourikas D, Pekarik V, Baeriswyl T, Grunditz A, Sadhu R, Nardo M. and Stoeckli ET (2005) Sonic hedgehog guides commissural axons along the longitudinal axis of the spinal cord. Nature Neurosci 8: 297-304.

Cafiero C, Marangi G, Orteschi D, Ali M, Asaro A, Ponzi E, Moncada A, Ricciardi S, Murdolo M, Mancano G, Contaldo I, Leuzzi V, Battaglia D, Mercuri E, Slavotinek AM, and Zollino M. (2015) Novel de novo heterozygous loss-of-function variants in MED13L and further delineation of the MED13L haploinsufficiency syndrome. Eur J Hum Genet. 23:1499-504. doi: 10.1038/ejhg.2015.19.

Calloni GW, Glavieux-Pardanaud C, Le Douarin NM, and Dupin E (2007) Sonic Hedgehog promotes the development of multipotent neural crest progenitors endowed with both mesenchymal and neural potentials. Proc Natl Acad Sci U S A 104: 19879-19884.

Davis MA, Larimore EA, Fissel BM, Swanger J, Taatjes DJ, and Clurman BE (2013) The SCF-Fbw7 ubiquitin ligase degrades MED13 and MED13L and regulates CDK8 module association with mediator. Genes Dev 27: 151–156.

De La Cruz López MV, González Criado M. (2011) Adaptación española del inventario de Desarrollo Battelle. Madrid: TEA

Delloye-Bourgeois C, Jacquier A, Charoy C, Reynaud F, Nawabi H, Thoinet K, Kindbeiter K, Yoshida Y, Zagar Y, Kong Y, Jones YE, Falk J, Chédotal A, and Castellani V (2015) PlexinA1 is a new Slit receptor and mediates axon guidance function of Slit C-terminal fragments. Nat Neurosci 1: 36-45

Fernell E, Olsson VA, Karlgren-Leitner C, Norlin B, Hagberg B and Gillberg C (1999) Autistic disorders in children with CHARGE association. Dev Med Child Neurol 41: 270-2.

Galbraith MD, Allen MA, Bensard CL, Wang X, Schwinn MK, Qin B, Long HW, Daniels DL, Hahn WC, Dowell RD, and Espinosa JM (2013) HIF1A employs CDK8-mediator to stimulate RNAPII elongation in response to hypoxia. Cell 153:1327-39.

Gilissen C, Hehir-Kwa JY, Thung DT, van de Vorst M, van Bon BW, Willemsen MH, Kwint M, Janssen IM, Hoischen A, Schenck A, Leach R, Klein R, Tearle R, Bo T, Pfundt R, Yntema HG, de Vries BB, Kleefstra T, Brunner HG, Vissers LE, and Veltman JA. (2014) Genome sequencing identifies major causes of severe intellectual disability. Nature 511: 344–347.

Guion-Almeida, ML and Richieri-Costa A (2009) Frontonasal dysplasia, severe neuropsychological delay, and midline central nervous system anomalies: report of 10 Brazilian male patients. Am J Med Genet A 149: 1006-1011.

Hennekam RC (2006) Rubinstein-Taybi syndrome. Eur J Hum Genet 14: 981-5.

Iossifov I, Ronemus M, Levy D, Wang Z, Hakker I, Rosenbaum J, Yamrom B, Lee YH, Narzisi G, Leotta A, Kendall J, Grabowska E, Ma B, Marks S, Rodgers L, Stepansky A, Troge J, Andrews P, Bekritsky M, Pradhan K, Ghiban E, Kramer M, Parla J, Demeter R, Fulton LL, Fulton RS, Magrini VJ, Ye K, Darnell JC, Darnell RB, Mardis ER, Wilson RK, Schatz MC, McCombie WR, and Wigler M. (2012) De novo gene disruptions in children on the autistic spectrum. Neuron 74:285-99. doi: 10.1016/j.neuron.2012.04.009.

Krebs AR, Demmers J, Karmodiya K, Chang NC, Chang AC, and Tora L (2010) ATAC and Mediator coactivators form a stable complex and regulate a set of non-coding RNA genes. EMBO Rep. 11: 541-7.

López-Ornat S, Gallego C, Gallo P, Karousou A, Mariscal S, and Martínez M (2005) MacArthur: Inventario de desarrollo comunicativo. Madrid: TEA.

Muncke N, Jung C, Rüdiger H Ulmer H, Roeth R, Hubert A, Goldmuntz E, Driscoll D, Goodship J, Schön K, and Rappold G. (2003) Missense variants and gene interruption in PROSIT240, a novel TRAP240-like gene, in patients with congenital heart defect (transposition of the great arteries). Circulation 108: 2843–2850.

Najmabadi H, Hu H, Garshasbi M Zemojtel T, Abedini SS, Chen W, Hosseini M, Behjati F, Haas S, Jamali P, Zecha A, Mohseni M, Püttmann L, Vahid LN, Jensen C, Moheb LA, Bienek M, Larti F, Mueller I, Weissmann R, Darvish H, Wrogemann K, Hadavi V, Lipkowitz B, Esmaeeli-Nieh S, Wieczorek D, Kariminejad R, Firouzabadi SG, Cohen M, Fattahi Z, Rost I, Mojahedi F, Hertzberg C, Dehghan A, Rajab A, Banavandi MJ, Hoffer J, Falah M, Musante L, Kalscheuer V, Ullmann R, Kuss AW, Tzschach A, Kahrizi K, and Ropers HH (2011) Deep sequencing reveals 50 novel genes for recessive cognitive disorders. Nature 478: 57–63.

Robins DL, Fein D, Barton ML, Green JA. (2001) The modified checklist for autism in toddlers: An initial study investigating the early detection of autism and pervasive developmental disorders. J Autism Dev Disord 31: 131-144.

Shashi V, Pena LD, Kim K, Burton B, Hempel M, Schoch K, Walkiewicz M, McLaughlin HM, Cho M, Stong N, Hickey SE, Shuss CM; Undiagnosed Diseases Network, Freemark MS, Bellet JS, Keels MA, Bonner MJ, El-Dairi M, Butler M, Kranz PG, Stumpel CT, Klinkenberg S, Oberndorff K, Alawi M, Santer R, Petrovski S, Kuismin O, Korpi-Heikkilä S, Pietilainen O, Aarno P, Kurki MI, Hoischen A, Need AC, Goldstein DB, Kortüm F. (2016) De novo truncating variants in ASXL2 are associated with a unique and recognizable clinical phenotype. Am J Hum Genet. 99:991-999.

Salie R, Niederkofler V, and Arber S (2005) Patterning molecules: multitasking in the nervous system. Neuron 45: 189-192.

Sener RN (1995) Rubinstein-Taybi syndrome: cranial MR imaging findings. Comput Med Imaging Graph 19: 417-8.

Utami KH, Winata CL, Hillmer AM, Aksoy I, Long HT, Liany H, Chew EG, Mathavan S, Tay SK, Korzh V, Sarda P, Davila S, and Cacheux V. (2014) Impaired development of neural-crest cell-derived organs and intellectual disability caused by MED13L haploinsufficiency. Hum Mutat. 35:1311-20. doi: 10.1002/humu.22636.

Valor LM, Pulopulos MM, Jiménez-Minchán M, Olivares R, Lutz B, and Barco A. (2011) Ablation of CBP in forebrain principal neurons causes modest memory and transcriptional defects and a dramatic reduction of histone acetylation but does not affect cell viability. J Neurosci 31: 1652-63.

Verpoorten R, Noens I, and van Berckelaer-Onnes I (2014) Manual ComFor: precursores de la comunicación. Ávila: Autismo Ávila.

Viosca J, López-Atalaya JP, Olivares R, Eckner R and Barco A (2010) Syndromic features and mild cognitive impairment in mice with genetic reduction on p300 activity: Differential contribution of p300 and CBP to Rubinstein-Taybi syndrome etiology. Neurobiol Dis 37:186-94.

Wada N, Javidan Y, Nelson S, Carney TJ, Kelsh RN, and Schilling TF (2005) Hedgehog signaling is required for cranial neural crest morphogenesis and chondrogenesis at the midline in the zebrafish skull. Development 132: 3977-3988.

Wang J, Weaver IC, Gauthier-Fisher A, Wang H, He L, Yeomans J, Wondisford F, Kaplan DR, and Miller FD (2010) CBP histone acetyltransferase activity regulates embryonic neural differentiation in the normal and Rubinstein-Taybi syndrome brain. Dev Cell 18: 114-25.

